# Improved adenine-HPLC method for quantifying yeast based on cellular DNA content

**DOI:** 10.64898/2026.03.13.711611

**Authors:** Yoshifumi Ohyama, Mamiko Shimamura, Yuki Asami, Dieter M. Tourlousse, Naoyuki Togawa, Koji Narita, Naoki Hayashi, Jun Terauchi, Yuji Sekiguchi, Hiroko Kawasaki, Takamasa Miura

## Abstract

Accurate quantification of fungi is important for a myriad of applications but remains challenging. Previously, we demonstrated that an approach called the adenine-HPLC method can quantify bacteria, including those with aggregating properties that are difficult to quantify using conventional methods, by measuring cellular adenine derived from DNA and converting the adenine amount to genome copy number, without being influenced by cell morphology. However, in this study, when this adenine-HPLC method was applied to the quantification of budding yeast as a model fungus, accurate measurement proved impossible. This limitation was attributed to adenine release from other adenine-containing biomolecules, such as RNA and ATP, and we therefore developed a method that suppresses adenine release from these molecules. This method involves reducing the temperature of the acid treatment and prewashing the cells before acid treatment. In addition, we incorporated a process that corrects for the naturally occurring free adenine level as background during total adenine measurement. The improved adenine-HPLC method based on these modifications enables accurate quantification of budding yeast using genomic DNA content in whole cells as the quantification unit.

## Introduction

Fungi are found in almost all environments on Earth and form the basis of material cycling in ecosystems by decomposing organic matter [1]. Fungi are also members of the human microbiota and are intimately involved in human health and disease [2–4]. Furthermore, beyond their historical contributions to food production and dietary culture through fermentation and brewing [5], fungi are widely utilized in modern industries to produce valuable substances such as enzymes, antibiotics, novel biomaterials, and biofuels [6], to improve agricultural productivity through soil amelioration [7], and to address environmental challenges through bioremediation and forest restoration [8, 9]. It is expected that diverse medical, industrial, and societal issues will be resolved through improved understanding, utilization, and modification of fungal functions related to metabolism, physiology, and ecology. In all cases, precise handling, including accurate quantification of fungi in laboratory and industrial settings, is fundamentally important.

Various techniques for quantifying fungi have been developed to date. For targeting a limited range of fungi, biochemical markers such as ergosterol and specific phospholipids have been utilized, and antibodies for ELISA have been developed [10, 11]. For more universal measurements, conventional colony counting by cultivation, microscopic observation, and flow cytometry have been used [12, 13]. In addition, techniques focusing on cellular DNA content, such as quantitative PCR and shotgun sequencing, have also been developed [14–16]. However, methods depending on visual or optical observations face difficulties due to unculturable traits, aggregation, unclear cell–cell boundaries, and multicellular structures. Moreover, methods focusing on DNA quantity face the challenge that differences in PCR efficiency caused by cross-species primer design for diverse fungal genome sequences compromise quantifiability [15, 17]. Additionally, dependence on DNA extraction methods to obtain DNA templates for PCR and sequencing introduces inaccuracies caused by differences in DNA extraction efficiency among fungi with diverse cell walls [18].

Many of the above problems are also commonly encountered in bacterial quantification [19–21]. We previously proposed another universal quantification method for bacteria referred to as the adenine-HPLC method [22–24]. This method quantifies bacteria based on genomic DNA copy number as the quantification unit and does not require a DNA extraction process, resulting in quantification that is independent of differences in cell morphology and cell wall structure. It should be noted that the adenine-HPLC method is designed for accurate quantification of a single bacterial species (that is, it is not suitable for measuring the individual quantities within mixtures of multiple species) and requires genome sequence information for quantification. In this method, bacteria are treated with acid to induce DNA depurination. The liberated adenine is then quantified by HPLC. This adenine quantity is combined with separately obtained genomic information to calculate the genomic DNA copy number. We have demonstrated that the adenine-HPLC method shows higher accuracy than microscopy and flow cytometry and provides equivalent accuracy across dozens of bacterial species [24]. We considered that if the adenine-HPLC method could be applied to fungi, it could become a new versatile quantitative method that could replace or complement previously developed techniques. In this study, we used experimental BY series strains [25] of unicellular budding yeast as a fungal model and confirmed that the adenine-HPLC method is not suitable for fungal quantification. Therefore, we sought to improve the method and succeeded in quantifying the genomic DNA copy number of budding yeast using a modified approach.

## Materials and Methods

### Genetic background and culturing conditions

Supplemental Table S1 lists the genomic information of *Escherichia coli* and the genotypes of the *Saccharomyces cerevisiae* strains BY4741, BY4742, and BY4743 (BY series strains). *E*. coli was cultured in 802 medium (https://www.nite.go.jp/nbrc/catalogue/NBRCMediumDetailServlet?NO=802) at 30 °C with rotation at 180 rpm for 20 h. The BY series strains were cultured in YPD or SD medium at 30 °C with rotation at 115 rpm for 24 h. YPD and SD media were prepared as described by Dunham *et al*[26]. SD medium was supplemented with nutrients at standard concentrations [26] according to the auxotrophic requirements of the BY series strains. Cultures were collected by centrifugation (4,000 × g, 4 °C, 10 min), washed once with cold phosphate-buffered saline (PBS), and centrifuged again to obtain cell pellets. The pellets were suspended in PBS containing 15% (v/v) glycerol (PBS–glycerol) and stored at −80 °C until use.

### Microscopic observation

Microscopic observation of *E. coli* was performed as described by Ohyama *et al*. [24]. For the BY series strains, cell stocks were diluted 50- to 100-fold with PBS–glycerol and used for bright-field microscopy with a Thoma cell counter (WATSON) according to the manufacturer’s instructions. Microscopic images were captured using ZEN software integrated into the ZEISS microscope system and analyzed using Fiji [27] to determine cell numbers. At least three independently cultured samples were used for microscopic counting, and at least 15 images were captured for each observation.

### DNA extraction

DNA from *E. coli* was extracted as described by Ohyama *et al*. [24]. To extract DNA from *S. cerevisiae*, stored cells were washed twice with wash buffer (50 mM Tris-HCl [pH 7.5], 50 mM EDTA [pH 8.0], 500 mM sorbitol, 50 mM 2-mercaptoethanol) and pelleted by centrifugation. The washed cells were suspended in Zymolyase solution (0.76 M sorbitol, 0.18 mg/mL Zymolyase^TM^-100T; Nacalai Tesque), incubated at 32 °C for 30 min, and pelleted by centrifugation. The pellet was then resuspended in lysis buffer (50 mM Tris-HCl [pH 7.5], 100 mM EDTA [pH 8.0], 200 mM NaCl, 0.5% [w/v] SDS, 0.4 mg/mL Proteinase K) and incubated at 56 °C for 20 min. The solution was then extracted with phenol/chloroform/isoamyl alcohol (25:24:1; PCI), and the aqueous phase was transferred to a new tube. An equal volume of 2-propanol was added, and the mixture was centrifuged to precipitate nucleic acids. After washing the pellet with 70% ethanol, RNase A solution (50 mM Tris-Cl [pH 8.0], 10 mM EDTA, 0.4 mg/mL RNase A) was added, and the mixture was incubated at 37 °C for 30 min. Serial PCI and chloroform extractions were performed on the RNase A-treated solution, and the supernatant was transferred to a new tube. Subsequently, 1/10 volume of 3 M sodium acetate and 2.5 volumes of ethanol were added, mixed, and centrifuged to precipitate nucleic acids. The nucleic acid pellet was resuspended in RNase A solution and incubated at 37 °C for 30 min. DNA was then purified from the RNase A-treated solution using the MagAttract HMW DNA Kit (QIAGEN) as follows. Two hundred μL of Buffer AL, 15 μL of MagAttract beads, and 280 μL of Buffer MB were added to the RNase A-treated solution and mixed thoroughly. The resulting DNA–bead complexes were captured using a magnetic stand and washed with the kit wash buffers according to the manufacturer’s instructions. Finally, DNA was eluted from the beads using Buffer AE. Serial PCI and chloroform extractions were performed on the eluate. The supernatant was transferred to a new tube, and 1/10 volume of 3 M sodium acetate and 2.5 volumes of ethanol were added and mixed to precipitate DNA. After centrifugation, the DNA pellet was washed with 70% ethanol and dissolved in Buffer EB (QIAGEN). The DNA concentration was measured using the Ouant-iT^TM^ PicoGreen^TM^ dsDNA Assay Kit (Thermo Fisher Scientific) and a microplate reader (Varioskan; Thermo Fisher Scientific) according to the manufacturer’s instructions.

### Sample treatment for measuring adenine content by HPLC and calculation of genomic DNA copy number from adenine content

Sample treatment using the adenine-HPLC method was conducted as previously described [22–24]. In this study, several modifications were incorporated into the adenine-HPLC method, as described below.

#### > Prewashing cells with SDS solution

A defined volume of PBS–glycerol cell suspension of the BY series strains was transferred to a new tube and adjusted to 1 mL with prewashing solution containing 0.2% (w/v) SDS. The suspension was incubated at 30 °C for 5 min with shaking at 500 rpm (ThermoMixer; Eppendorf). After centrifugation (6,000 × g, 25 °C, 1 min), the supernatant was removed by pipetting. The pellet was then adjusted to 120 μL with solution containing 0.1% (w/v) SDS (treatment solution) and mixed thoroughly by pipetting. To determine the optimal SDS concentration in the prewashing solution, 0%, 0.05%, 0.1%, 0.2%, or 0.5% (w/v) SDS was tested (Supplemental Fig. S3).

#### > Acid treatment

To 120 μL of the cells in treatment solution, 30 μL of 1 N HCl was added and mixed. The mixture was incubated at 35 °C with shaking at 500 rpm (ThermoMixer; Eppendorf) for the times indicated in the Results section. Subsequently, 50 μL of 1 N NaOH was added, and the tube was transferred to a preheated 100 °C block incubator and incubated for 30 min with occasional tapping. The tube was centrifuged (20,000 × g, 25 °C, 5 min), and 150 μL of the supernatant was transferred to a new tube. To neutralize the solution, 15 μL of 1 N HCl and 60 μL of 0.4 M ADA buffer (pH 6.7; FUJIFILM) were added and mixed thoroughly by vortexing. For acid treatment of purified DNA (Fig. 1) and adenine-containing biomolecules of interest (Fig. 2 and Fig. S1), 1.5 μg of purified DNA, 8.1 μg of tRNA and poly(A) RNA, or 12 nmol of ATP and AMP was used per reaction. Acid treatment was performed as described above, except that the total reaction volume was reduced to one-third. When unwashed cells were used, 120 µL of cells in treatment solution was prepared directly from a defined volume of cell stock and used as the starting material.

**Fig. 1.**
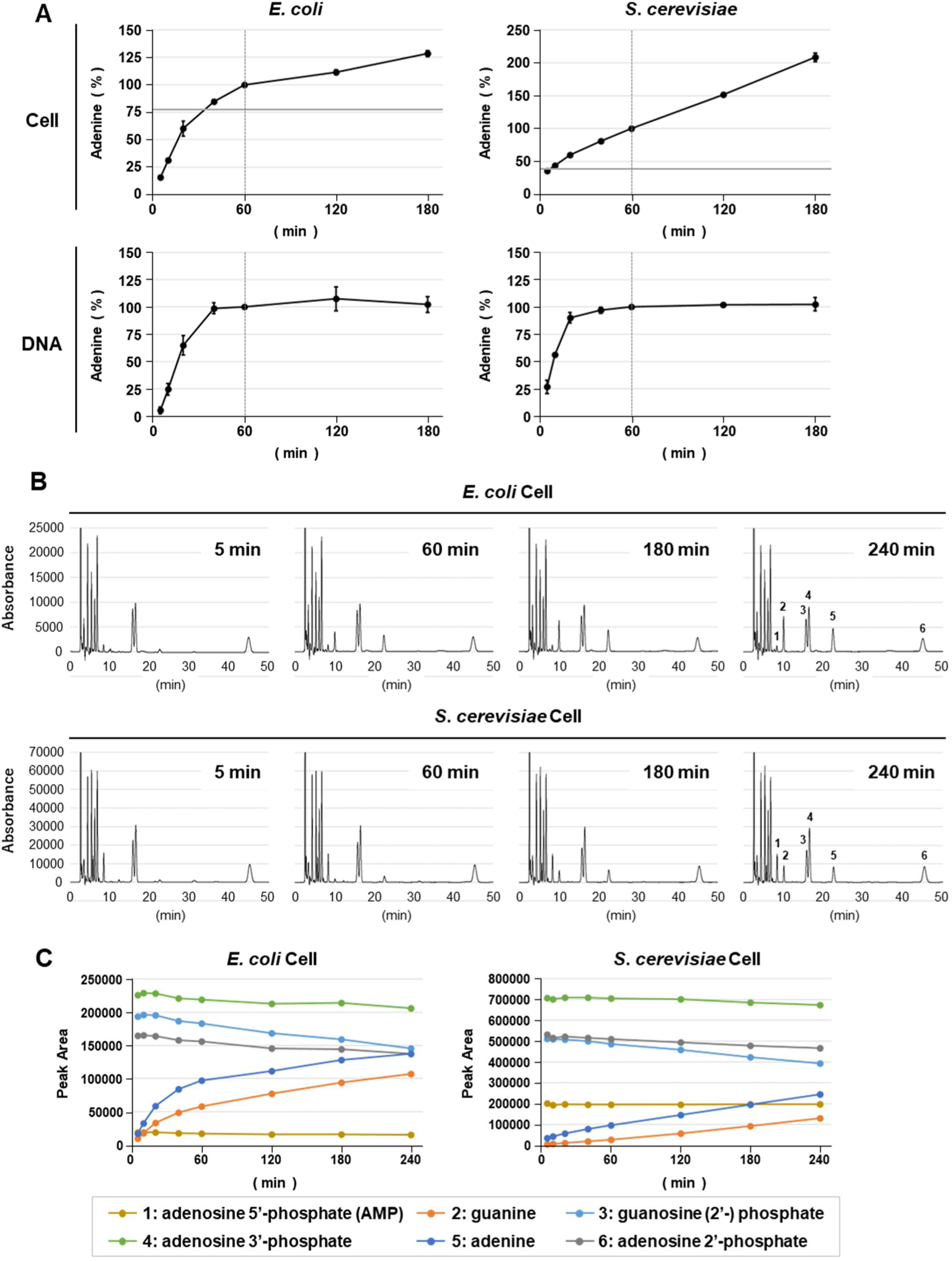
Adenine release kinetics in *E*. *coli* and *S*. *cerevisiae* determined by the adenine-HPLC method. (A) Using *E*. *coli* NBRC 3301 and *S*. *cerevisiae* BY4742 cells or their purified DNA, adenine release was measured at different time points during acid treatment (5, 10, 20, 40, 60, 120, and 180 min; x-axis) by the adenine-HPLC method. The measured adenine amount is plotted on the y-axis as a relative value normalized to the 60-min time point (100%). The vertical dashed line in the cell graph indicates the 60-min time point used for acid treatment in the original adenine-HPLC method. The gray horizontal line indicates the relative ratio of adenine estimated from microscopic cell counts (assuming a haploid genome per cell) to the adenine amount at 60 min determined by the adenine-HPLC method. Each measurement was performed in triplicate using independently prepared samples. (B) Representative chromatograms obtained at each acid treatment time point in the adenine-HPLC method are shown. Each peak is assigned a number corresponding to the following molecular species: 1, adenosine 5′-phosphate (AMP); 2, guanine; 3, guanosine (2′-) phosphate; 4, adenosine 3′-phosphate; 5, adenine; and 6, adenosine 2′-phosphate. (C) Peak areas of the molecules shown in (B) are plotted across all time points (5, 10, 20, 40, 60, 120, 180, and 240 min). A representative result is shown (n = 2).

**Fig. 2.**
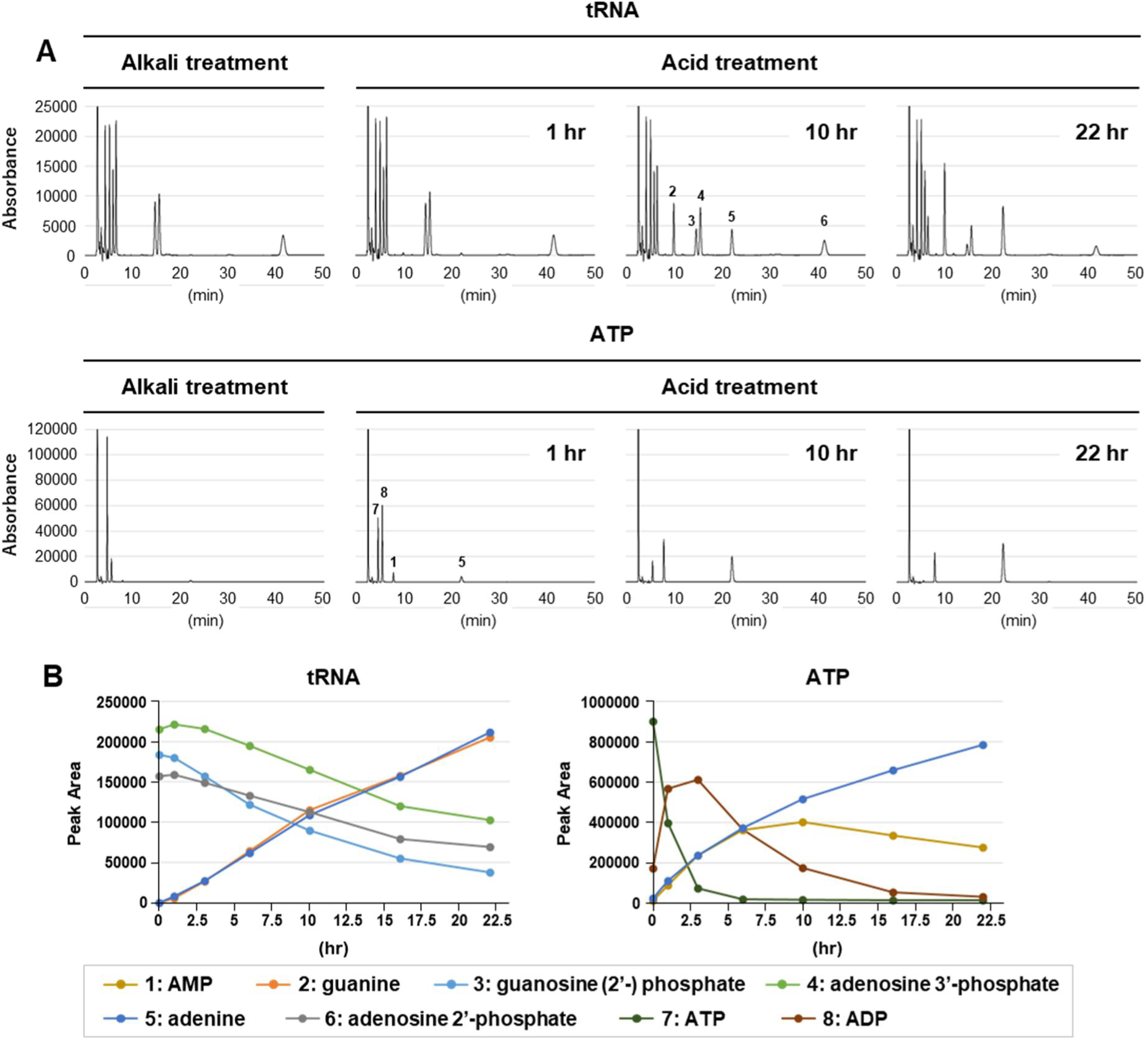
Adenine release kinetics of tRNA and ATP. (A) Representative chromatograms of tRNA and ATP subjected to alkali treatment or acid treatment at 60 °C for the indicated time points are shown. Each peak is assigned a number corresponding to the molecular species described in Fig. 1 and as follows: 7, adenosine 5′-triphosphate (ATP); and 8, adenosine 5′-diphosphate (ADP). (B) Peak areas of the molecules shown in (A) are plotted across all time points (1, 3, 6, 10, 16, and 22 h). A representative result is shown (n = 2).

#### > Alkali treatment

For alkaline treatment, the same stock solution of cells or compounds was used at the same volume as in the acid treatment. To 120 μL of cell suspension prepared as described above, 30 μL of 1 N HCl and 50 μL of 1 N NaOH were added simultaneously. The mixture was transferred to a preheated 100 °C block incubator and incubated for 30 min with occasional tapping. The tube was centrifuged (20,000 × g, 25 °C, 5 min), and 150 μL of the supernatant was transferred to a new tube. To neutralize the solution, 15 μL of 1 N HCl and 60 µL of 0.4 M ADA buffer were added and mixed thoroughly by vortexing. For purified DNA or biomolecules, the alkaline treatment was performed as described above, with the reaction volume reduced to one-third.

#### > HPLC measurement and calculation of genomic DNA quantity

Prior to HPLC measurement, the acid- or alkali-treated solution was thoroughly vortexed and centrifuged (20,000 × g, 25 °C, 2 min) to precipitate insoluble materials. The supernatant was applied to a Nexera-i LC-2040C system (Shimadzu), and measurement was conducted as follows. Ten μL of sample was injected into a reverse-phase column (Inertsil ODS-3; 5 μm particle size, 4.6 × 250 mm; GL Sciences) and pumped at a flow rate of 1 mL/min using an isocratic solvent buffer (30 mM ammonium acetate, 1 mM CyDTA, 10 mM NaH_2_PO_4_, 2% [v/v] methanol; pH 6.3). The analyte signal was detected at 260 nm for 50 min. The area under the curve (AUC) of each peak was calculated using LabSolutions™ software (Shimadzu), and the AUC of the alkali-treated sample was subtracted from that of the acid-treated sample. The resulting value was converted to adenine quantity using a standard curve generated as described by de Bruin and Birnbiom [28]. Conversion of adenine quantity to genomic DNA copy number was performed using the Excel sheet provided in Supplemental Table S2.

## Results

### Applicability of the adenine-HPLC method to budding yeast

We evaluated the applicability of the adenine-HPLC method [22–24] to *S. cerevisiae* BY series strains (BY4741 and BY4742 are haploids representing a mating-type pair, and BY4743 is their zygotic diploid). Because the adenine-HPLC method quantifies genomic DNA copy number, we first used the haploid BY4742 strain to facilitate interpretation of the relationship between genomic DNA content and cell number. In addition, *E. coli* was used as a control, for which the quantitative accuracy of the adenine-HPLC method has already been validated. In the original report [28], adenine release from DNA was observed under acidic conditions (0.2 N HCl) at 60 °C over time, and 100% of adenine was released within 60 min of incubation, as indicated by a plateau in the adenine release curve. However, the kinetics of adenine release from bacterial cells had not been investigated. Therefore, we first examined the kinetics of adenine release from *E. coli* and BY4742 cells alongside their purified DNA (Fig. 1). While the adenine release rate in *E. coli* cells declined markedly up to 120 min, similar to the kinetics observed for purified DNA, it continued to increase proportionally with incubation time in BY4742 (Fig. 1A and 1C). These results indicate that an acid treatment time of 60 min is not adequate to quantify adenine in fungi. To investigate the underlying reason, we carefully examined the time-dependent changes in the HPLC chromatograms. As shown in Fig. 1B, in addition to the purine bases adenine and guanine, peaks corresponding to RNA-derived adenosine 2′- or 3′-phosphates and guanosine (2′-) phosphate were observed, consistent with a previous report [28]. In addition, a peak corresponding to adenosine 5′-phosphate (AMP), which was not derived from RNA (see next section), was detected. Quantification of these peaks (Fig. 1C) revealed that the levels of adenosine phosphates and guanosine phosphate gradually decreased at a slow rate in both *E. coli* and budding yeast. Moreover, the ratio of purine bases to adenosine and guanosine phosphates was markedly higher in BY4742 than in *E. coli* (Fig. 1C), indicating that BY4742 contains a greater amount of RNA and AMP than *E. coli*.

### Adenine release kinetics of RNA and ATP

We hypothesized that adenine release from abundant RNA contributes to the continuous increase in adenine release observed in BY4742 cells. In addition, ATP is a representative adenine-containing biomolecule. Because eukaryotic cells efficiently produce ATP in mitochondria, they are expected to contain substantial amounts of ATP. Therefore, we investigated the adenine release kinetics of purified RNA (tRNA and poly[A] RNA), as well as ATP and AMP. Fig. 2 shows the behavior of tRNA and ATP over time under acidic conditions (0.2 N HCl, 60 °C). In this experiment, we also conducted alkaline treatment alone to degrade biomolecules, particularly to monomerize RNA (see Materials and Methods), in order to measure the baseline levels of each molecule by HPLC without acid treatment. Notably, alkaline treatment is normally performed in the adenine-HPLC method after acid treatment to monitor monomerized RNA. As shown in Fig. 2, purine release from tRNA progressed in association with the time-dependent degradation of adenosine and guanosine phosphates during prolonged acid treatment at 60 °C. Similarly, adenine release from ATP was observed in association with stepwise degradation from ATP to ADP and from ADP to AMP (Fig. 2B). Adenine release from AMP was also observed, although at a slower rate than from ATP (Supplemental Fig. S1). Additionally, for poly(A) RNA, which was used as a substitute for the poly(A) tail of eukaryotic mRNA, adenine release accompanying poly(A) degradation was observed at a slower rate than for tRNA (Supplemental Fig. S1). These results indicate that RNA and ATP may serve as sources of the continuous increase in adenine observed following acid treatment in budding yeast. Furthermore, the results obtained for tRNA and poly(A) RNA demonstrate that the adenosine 5′-phosphate (AMP) detected in cells after acid treatment was not a degradation product of RNA.

### Suppression of adenine release from RNA and ATP

The above results suggested that inhibiting adenine release from RNA and ATP may enable more accurate determination of DNA-derived adenine levels. Therefore, we investigated which parameters of the original adenine-HPLC method could affect adenine release from RNA and ATP. We found that adenine release occurred in a temperature-dependent manner (data not shown). Furthermore, reducing the acid treatment temperature to 35 °C (compared with 60 °C in the experiments described above) nearly completely suppressed adenine release from tRNA and poly(A) RNA and, although less effectively, suppressed release from ATP and AMP (Fig. 3 and Supplemental Fig. S2).

**Fig. 3.**
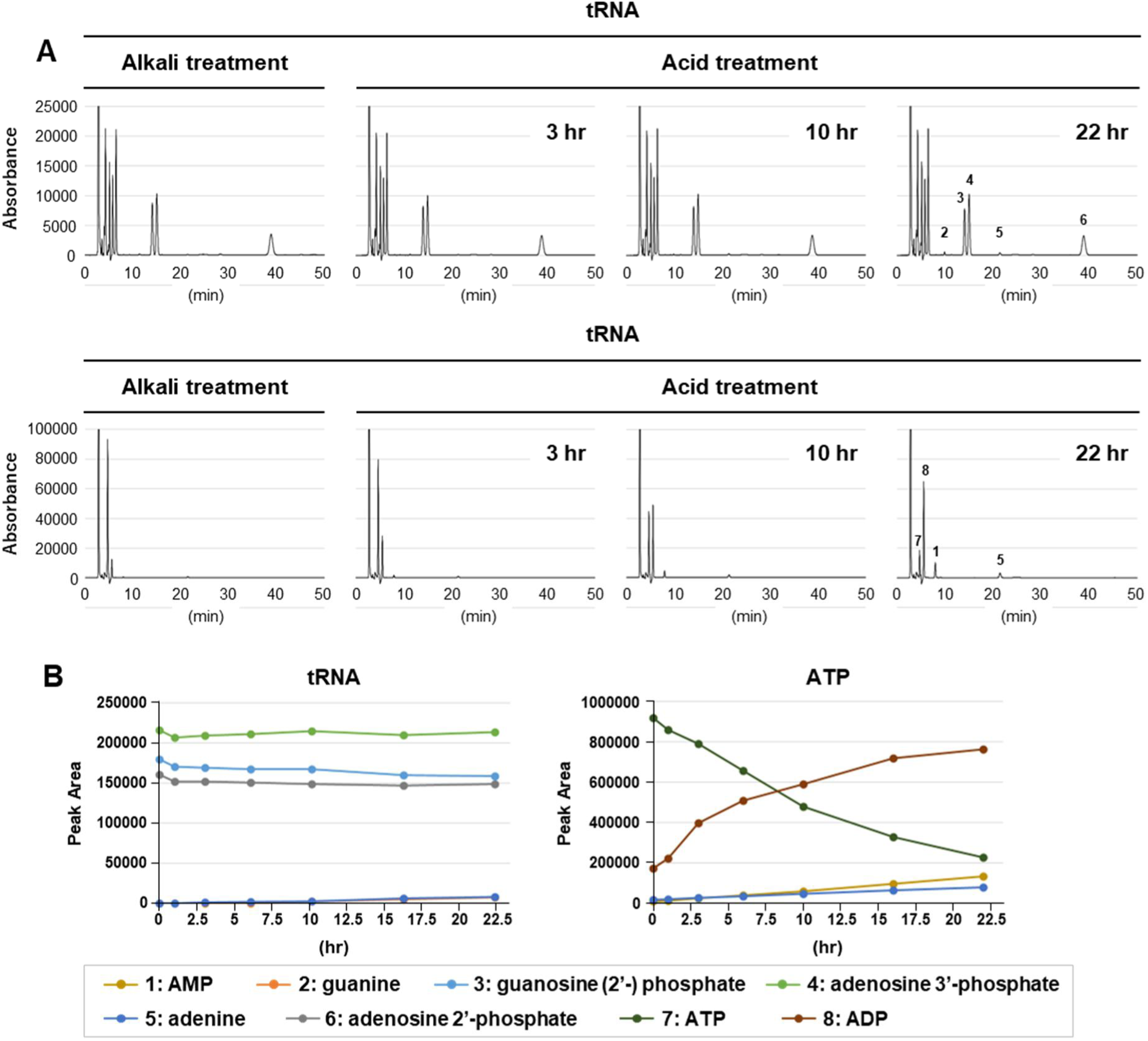
Suppression of adenine release from tRNA and ATP. (A) Representative chromatograms of tRNA and ATP subjected to alkali treatment or acid treatment at 35 °C for the indicated time points are shown. Each peak is assigned a number corresponding to the molecular species described in Fig. 1 and Fig. 2. (B) Peak areas of the molecules shown in (A) are plotted across all time points (1, 3, 6, 10, 16, and 22 h). A representative result is shown (n = 2).

### Suppression of adenine release from molecules other than RNA and ATP in cells

Although acid treatment at 35 °C reduced adenine release from RNA, ATP, and AMP, this does not necessarily imply that adenine release from other adenine-containing molecules in cells would also be suppressed under these conditions. It has been reported that the detergent SDS denatures cell membranes and permeabilizes cells, allowing cellular contents to diffuse into the extracellular space [29]. These effects were shown to depend on SDS concentration and treatment time [29]. Therefore, we hypothesized that controlling the stringency of SDS treatment might selectively remove small molecules, such as monomers, while retaining polymers within cells, given that most adenine-containing molecules are monomeric except for DNA and RNA. We investigated such conditions and found that incubation of BY4743 in SDS solution at 30 °C for 5 min (SDS washing) removed AMP, an indicator of intracellular monomers, in an SDS concentration-dependent manner while maintaining nearly constant RNA levels (Fig. S3). Furthermore, HPLC analysis of BY4743 with or without SDS washing followed by alkaline treatment showed that BY4743 contains endogenous adenine and guanine, which were also removed by SDS washing. In contrast, such endogenous purines were barely detectable in *E. coli* (Fig. 4B).

**Fig. 4.**
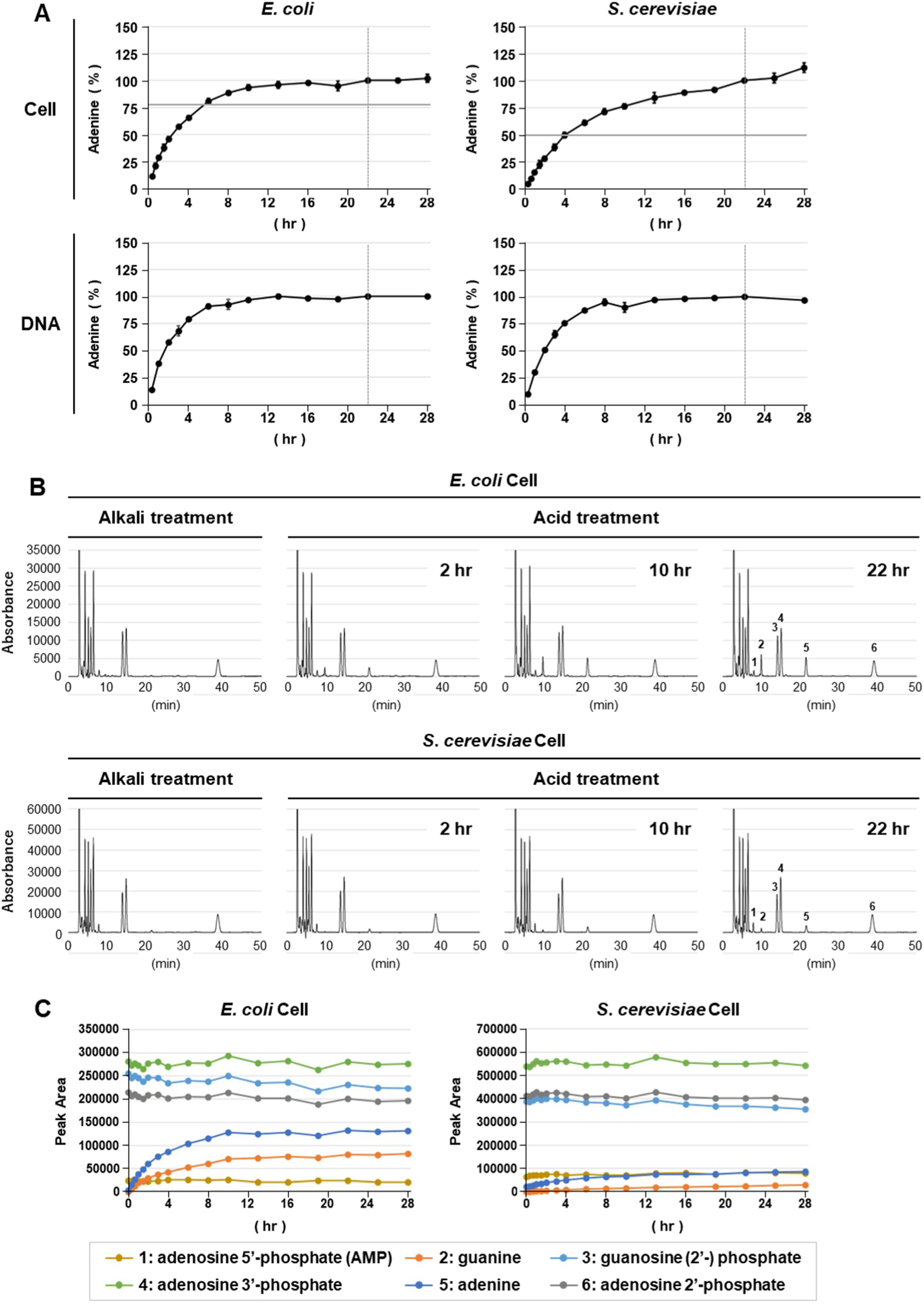
Adenine release kinetics in *E*. *coli* and *S*. *cerevisiae* under conditions that suppress adenine release from molecules other than DNA. (A) Alkali treatment and acid treatment at 35 °C were performed for the indicated time points using *E*. *coli* NBRC 3301 and *S*. *cerevisiae* BY4742 cells (0.3, 0.7, 1, 1.5, 2, 3, 4, 6, 8, 10, 13, 16, 19, 22, 25, and 28 h) or their purified DNA (0.3, 1, 2, 3, 4, 6, 8, 10, 13, 16, 19, 22, and 28 h). The measured adenine amount is shown relative to the 22-h time point (100%). The vertical dashed line and horizontal gray line are indicated as described in Fig. 1. Each measurement was performed three times using independently prepared samples. (B) Representative chromatograms obtained at selected acid treatment time points in (A) are shown. Each peak is assigned a number corresponding to the molecular species described in Fig. 1. (C) Peak areas of the molecules shown in (B) are plotted across all time points indicated in (A). A representative result is shown (n = 3).

### Adenine release kinetics under conditions that suppress adenine release from molecules other than DNA

Theoretically, prewashing cells with SDS before acid treatment at 35 °C should allow adenine release predominantly from DNA within cells. However, complete removal of small molecules, including AMP, without affecting RNA levels proved difficult. In many cases, SDS prewashing reduced small-molecule levels partially rather than eliminating them entirely. For example, washing BY4742 with 0.2% SDS at 30 °C for 5 min reduced AMP and endogenous adenine levels by approximately one-third and one-half, respectively (compare Fig. 4 and Fig. S4, alkali treatment data). Therefore, endogenous adenine was considered background relative to DNA-derived adenine. We measured endogenous adenine levels in alkali-treated samples and incorporated a step into the adenine-HPLC method in which this value was subtracted from the adenine content of the acid-treated sample. Using prewashed BY4742, unwashed *E. coli* (which was unsuitable for SDS washing, likely due to its thin cell wall), and their purified DNA, we monitored adenine release during acidic incubation at 35 °C over time (Fig. 4A). Up to 22 h, adenine release in BY4742 gradually approached a near-plateau phase. In contrast, adenine release in *E. coli* reached a plateau at 35 °C, consistent with the kinetics observed for its purified DNA. By comparing the adenine amount at 22 h with the amount expected from the same number of cells counted by microscopy, assuming a haploid genome per cell, we found that adenine levels at 22 h were approximately 2.0-fold higher in BY4742 and approximately 1.3-fold higher in *E. coli* than the levels estimated by microscopy (Fig. 4A). These observations may be partially explained by assuming that a proportion of cells were undergoing DNA replication (see Discussion).

### Importance of the SDS prewashing process

To determine whether SDS prewashing improved the quantitative accuracy for BY4742, we compared the adenine release kinetics of BY4742 cells with or without prewashing prior to acid treatment at 35 °C (Fig. S4). The adenine release curves showed a characteristic difference at 13 h (Fig. S4B). Specifically, before 13 h, the curves were nearly identical regardless of whether prewashing was performed; however, after 13 h, the rate of the continuous increase in adenine was lower in the washed cells. We infer that the continued increase in adenine after 13 h results from adenine release from adenine-containing molecules other than DNA, such as ATP. Because prewashing reduced this effect, we concluded that it is critical for more accurate quantification of adenine derived from BY4742 genomic DNA.

### Procedure of the new adenine-HPLC method

Based on the results described above, we modified the adenine-HPLC method as schematically illustrated in Fig. 5. First, cells are prewashed with SDS solution under optimized conditions. Next, alkaline treatment and acid treatment at 35 °C are performed independently. The adenine content of both alkali-treated and acid-treated samples is measured by HPLC, and the adenine content of the alkali-treated sample is subtracted from that of the acid-treated sample. This value is then converted to genomic DNA copy number using the procedure described in the Materials and Methods section. Hereafter, we refer to this modified approach as the new adenine-HPLC method. It should be noted that some cells, such as *E. coli*, are unsuitable for the SDS prewashing step; in such cases, this step may be omitted.

**Fig. 5.**
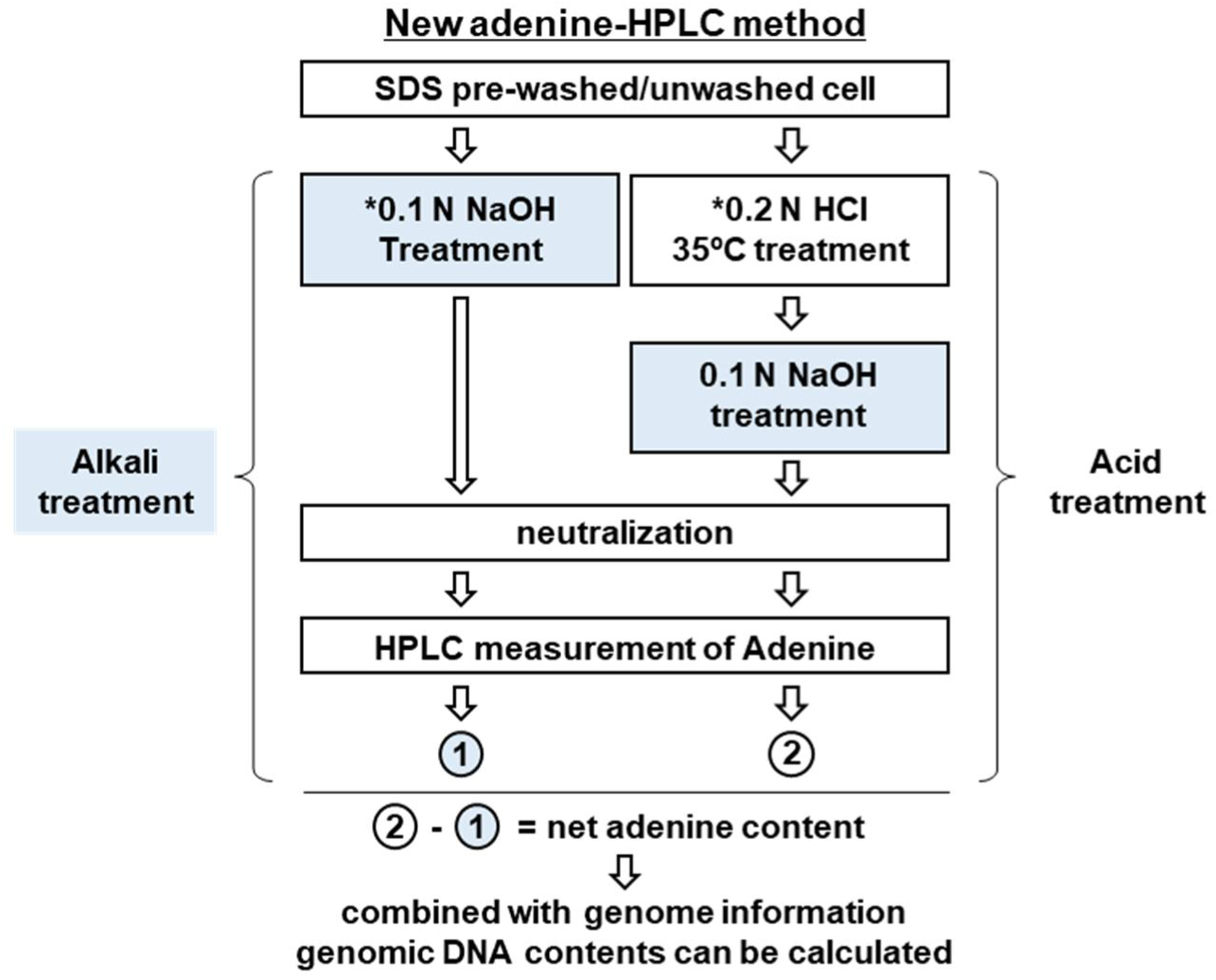
Schematic overview of the new adenine-HPLC method. The procedure of the new adenine-HPLC method is schematically illustrated. *The cell suspension containing 0.1% SDS was used for both alkali and acid treatment reactions (see Materials and Methods).

### Quantitative accuracy of the new adenine-HPLC method

To evaluate the quantitative accuracy of the new adenine-HPLC method for measuring budding yeast genomic DNA content, we utilized the ploidy differences among the BY series strains. We compared genomic DNA content among microscopically equivalent numbers of haploid BY4741 and BY4742 cells and diploid BY4743 cells (Fig. 6A), with or without SDS prewashing. If genomic DNA content was accurately measured by the new adenine-HPLC method, the same number of diploid and haploid cells would be expected to exhibit a twofold difference in genomic DNA content. As shown in Fig. 6B, genomic DNA content was approximately 1.9-fold higher in prewashed diploid cells than in haploid cells, indicating that the quantification closely matched the expected value. In contrast, cells without prewashing showed genomic DNA values approximately 15% higher than those of prewashed cells, consistent with the results shown in Supplemental Fig. S4B. Furthermore, the genomic DNA ratio between haploid and diploid cells in the unwashed group was 1.6-fold, indicating reduced quantitative accuracy. These results demonstrate that the new adenine-HPLC method provides sufficient quantitative accuracy to measure genomic DNA copy number in budding yeast.

**Fig. 6.**
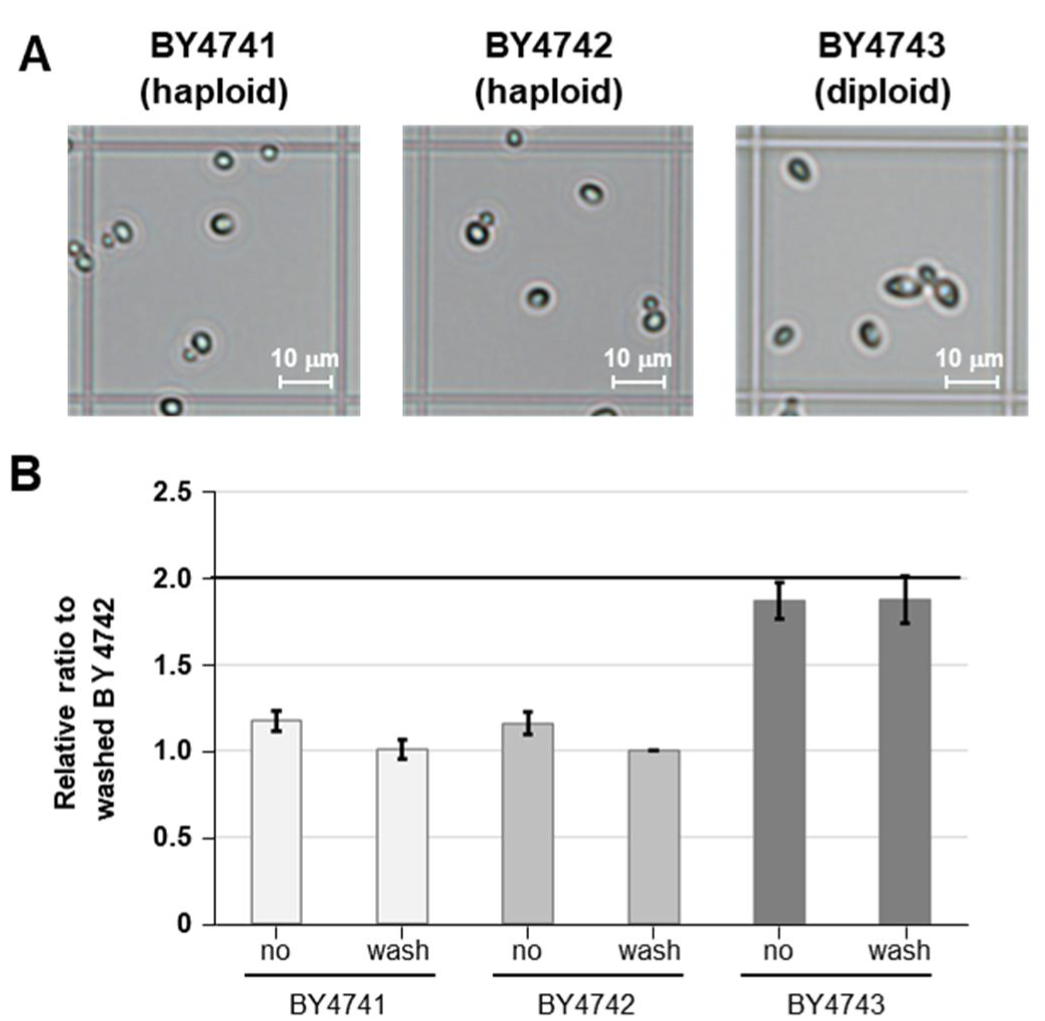
Validation of the quantitative accuracy of the new adenine-HPLC method. (A) Representative microscopic images of BY series strains on a hemocytometer are shown. (B) Genomic DNA copy number determined by the new adenine-HPLC method (22-h acid treatment) using equivalent cell numbers obtained from the measurements in (A). Results for each strain, with or without SDS prewashing, are shown. Genomic DNA copy number is expressed as a ratio relative to prewashed BY4742 cells.

## Discussion

Previously, we successfully quantified bacterial genomic DNA in whole cells directly using the adenine-HPLC method; however, this method was not applicable to budding yeast, prompting its modification. In budding yeast, in addition to adenine-containing biomolecules such as RNA and ATP, the intracellular level of free adenine is substantially higher than that in bacteria. We found that suppressing adenine release from biomolecules other than DNA by lowering the acid-treatment temperature, performing SDS prewashing, and subtracting endogenous adenine levels enabled selective measurement of DNA-derived adenine, thereby allowing calculation of genomic DNA copy number in budding yeast.

During these improvements, we determined that the addition of SDS during both alkali and acid treatments enhanced quantitative accuracy. Accordingly, SDS is employed in two distinct ways in the new adenine-HPLC method: to remove intracellular small molecules and to improve acid and alkali treatment reactions (Fig. 5 and Materials and Methods). As reported by An *et al.*, acid treatment of DNA produces viscous precipitates during the reaction, which complicates sample handling for HPLC measurement [30]. We found that formation of this viscous precipitate was almost completely suppressed by adding SDS to the acid treatment reaction. In addition, we observed that the rate of increase in the adenine release curve was faster in both *E*. *coli* DNA and cells than in reactions without SDS, indicating that SDS increased the adenine release rate (data not shown). This effect was not clearly observed in budding yeast and therefore cannot be explained by the prediction that nucleosomal DNA is more tolerant to acid-induced adenine release [30]. We speculate that this difference may reflect attenuation of the sequence dependence of adenine release rates reported for single-stranded DNA [30].

We compared the adenine amount determined by the new adenine-HPLC method with that estimated by microscopic counting (Fig. 4A). For *E*. *coli*, the adenine amount was approximately 1.3-fold relative to the microscopic cell count, falling within the expected range (1–2) when accounting for cells undergoing DNA replication within the measured population; this was therefore considered a reasonable result. In contrast, the ratio for budding yeast was approximately 2.0. We presume that this relatively higher value stems, at least in part, from the accumulation of adenine observed after 13 h of acid treatment, as shown in Fig. 4A and Supplemental Fig. S4A and S4B. Therefore, we reconsidered the adenine components that accumulated after 13 h of acid treatment.

Adenine release from DNA was rapid and was essentially completed within 13 h of acid treatment at 35 °C (Fig. 4A). In contrast, adenine release from ATP and AMP was slow and increased linearly at 35 °C (Fig. 3B and Fig. S2B). Under the current HPLC measurement conditions, ATP could not be quantified separately from other cellular components. However, as shown in Fig. 2B and Fig. 3B, ATP was converted to AMP following acid treatment. Therefore, changes in AMP levels after acid treatment may reflect intracellular ATP levels. Because budding yeast exhibited higher AMP levels than *E*. *coli* (Fig. 1C), we inferred that ATP was also present at higher levels in budding yeast. To examine this possibility, we simulated the adenine release reaction in BY4742 under the following assumptions. First, for adenine release from DNA, we used the adenine release curve of BY4742 purified DNA shown in Fig. 4A. Second, we used the values in Fig. 3B as the rate of adenine release from ATP at 35 °C. Third, we assumed that the ATP amount, expressed as the peak area ratio in the HPLC chromatogram, was either twofold or fourfold higher than the amount of adenine derived from DNA (in BY4742 shown in Fig. 4C and Fig. S4B, the AMP peak area relative to the maximal adenine peak area in the adenine release curve was similar in SDS-prewashed cells and approximately threefold higher in unwashed cells). When adenine release was assumed to originate solely from these two components in BY4742, the modeled adenine release curve was as shown in Supplemental Fig. S5. Although the absolute release rates differed, the simulated kinetics largely recapitulated the effect of SDS prewashing observed in BY4742 (Supplemental Fig. S4). These results support the hypothesis that ATP/ADP/AMP are major contributors to the gradual accumulation of adenine observed in BY4742 after 13 h of acid treatment. In contrast, in *E*. *coli*, the relatively low ATP levels likely explain why no apparent adenine accumulation was observed after 13 h under acid treatment at 35 °C (Fig. 4A).

In this study, we demonstrated the quantitative capability of the new adenine-HPLC method using budding yeast and *E*. *coli*. Beyond this evidence, we propose that this method is theoretically applicable to a broad range of microorganisms, provided that the SDS prewashing step is optimized. In addition, we believe that quantifying microbial abundance based on genomic DNA copy number is consistent with current and emerging trends in microbial research. High-throughput DNA sequencing technologies are now the primary tools for identifying microorganisms and investigating microbial composition and abundance in diverse environments. In these approaches, the obtained sequences, such as whole genomes and amplicons, serve as the fundamental units representing microorganisms. These sequence-based units are either equivalent to or proportional to genomic DNA copy number, which constitutes the fundamental unit in the new adenine-HPLC method, thereby indicating conceptual compatibility. Although this method requires genomic information to quantify genomic DNA copy number, technological advances are ongoing that enable more convenient and cost-effective determination of microbial genomic sequences [31]. We anticipate that the new adenine-HPLC method may contribute significantly to the future advancement of microbiology as a quantitative approach for measuring microbial abundance.

## Acknowledgments

We thank Mariko Akasaka for her helpful advice on manuscript preparation. We also thank Dr. Wakao Fukuda for technical assistance with the HPLC measurements.

## Supplemental Materials

**Fig. S1.**
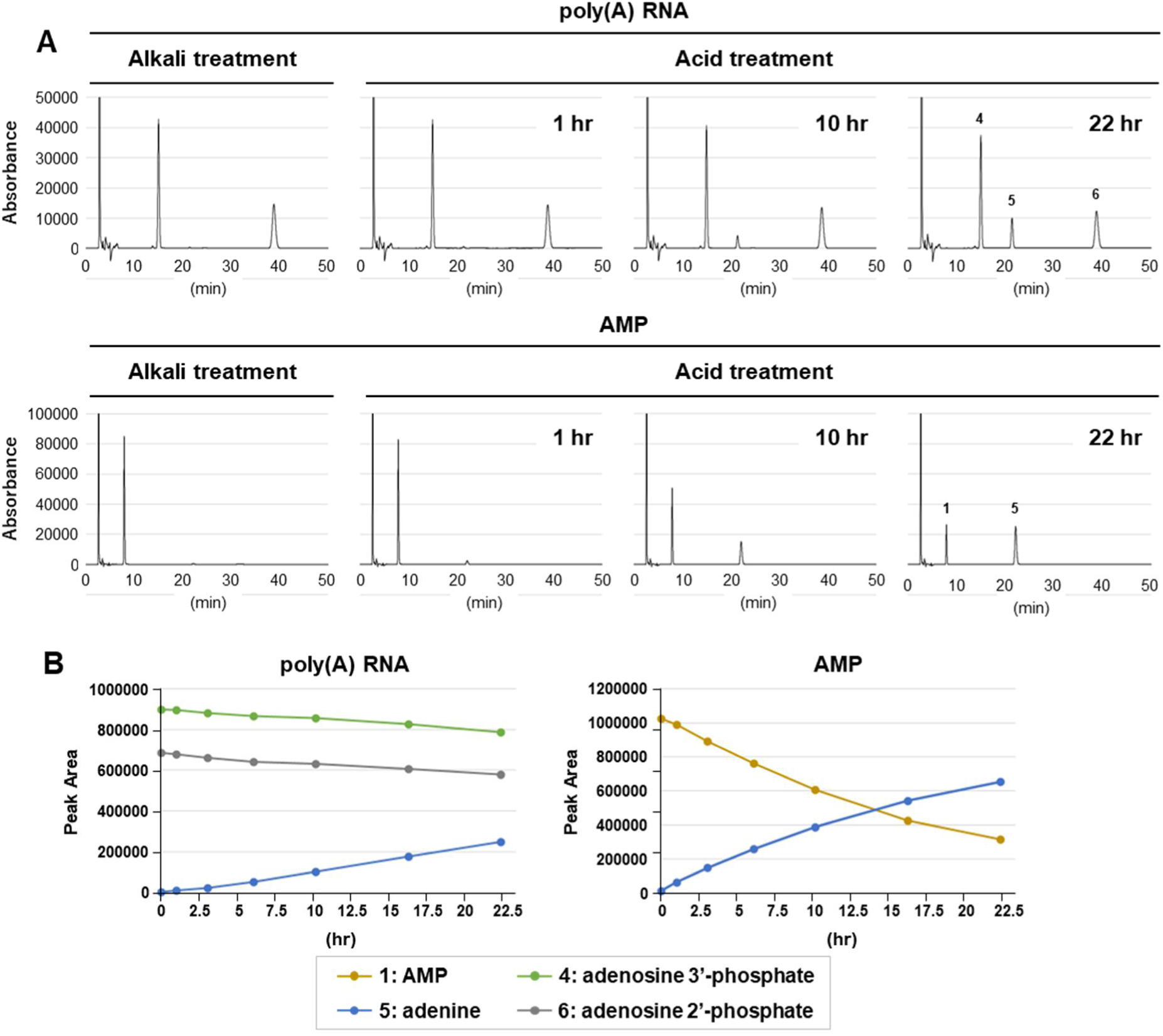
Adenine release kinetics of poly(A) RNA and AMP. (A) Representative chromatograms of poly(A) RNA and AMP subjected to alkali treatment or acid treatment at 60 °C for the indicated time points are shown. Each peak is assigned a number corresponding to the molecular species described in Fig. 1 and Fig. 2. (B) Peak areas of the molecules shown in (A) are plotted across all time points (1, 3, 6, 10, 16, and 22 h). A representative result is shown (n = 2).

**Fig. S2.**
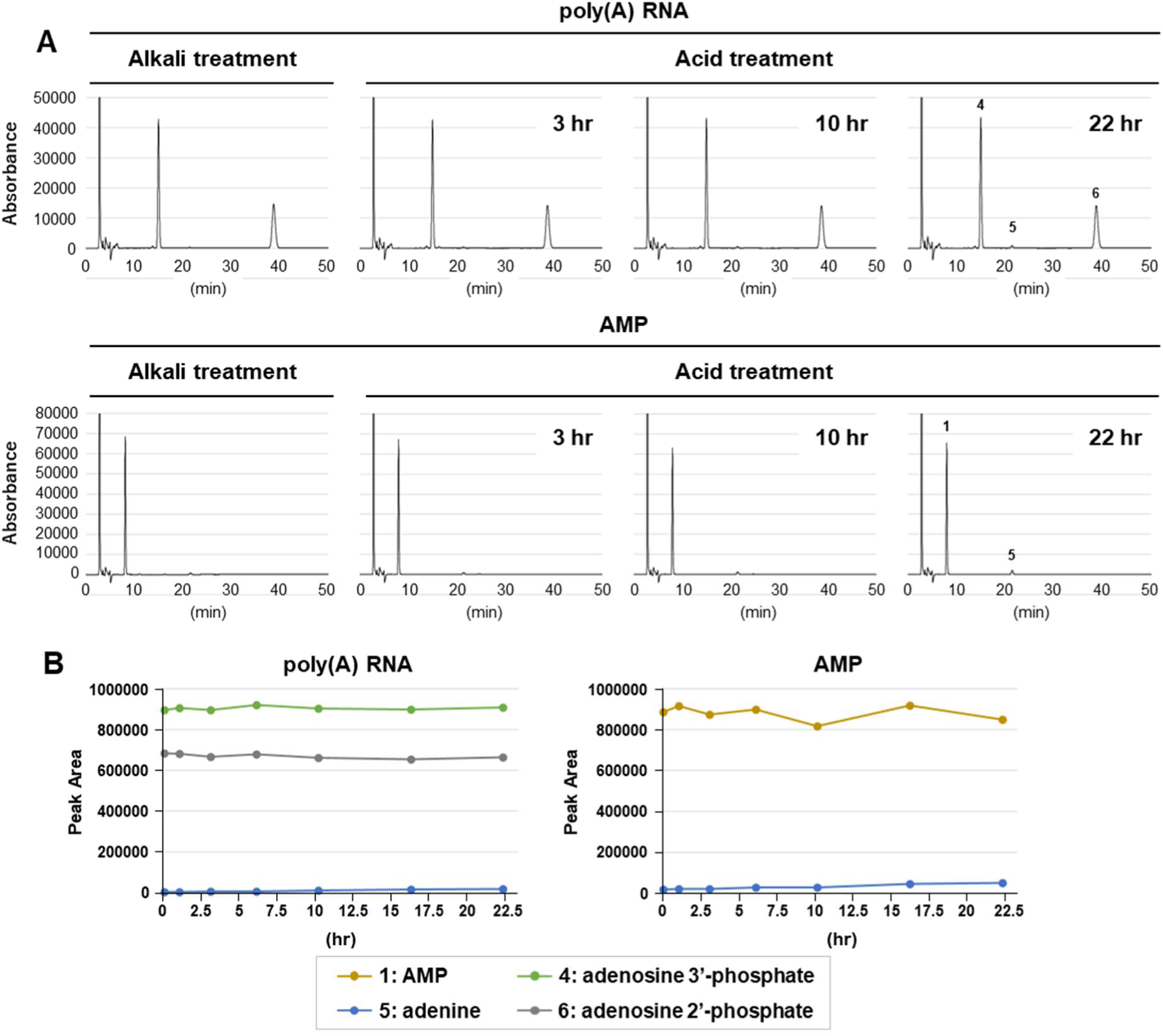
Suppression of adenine release from poly(A) RNA and AMP. (A) Representative chromatograms of poly(A) RNA and AMP subjected to alkali treatment or acid treatment at 35 °C for the indicated time points are shown. Each peak is assigned a number corresponding to the molecular species described in Fig. 1 and Fig. 2. (B) Peak areas of the molecules shown in (A) are plotted across all time points (1, 3, 6, 10, 16, and 22 h). A representative result is shown (n = 2).

**Fig. S3.**
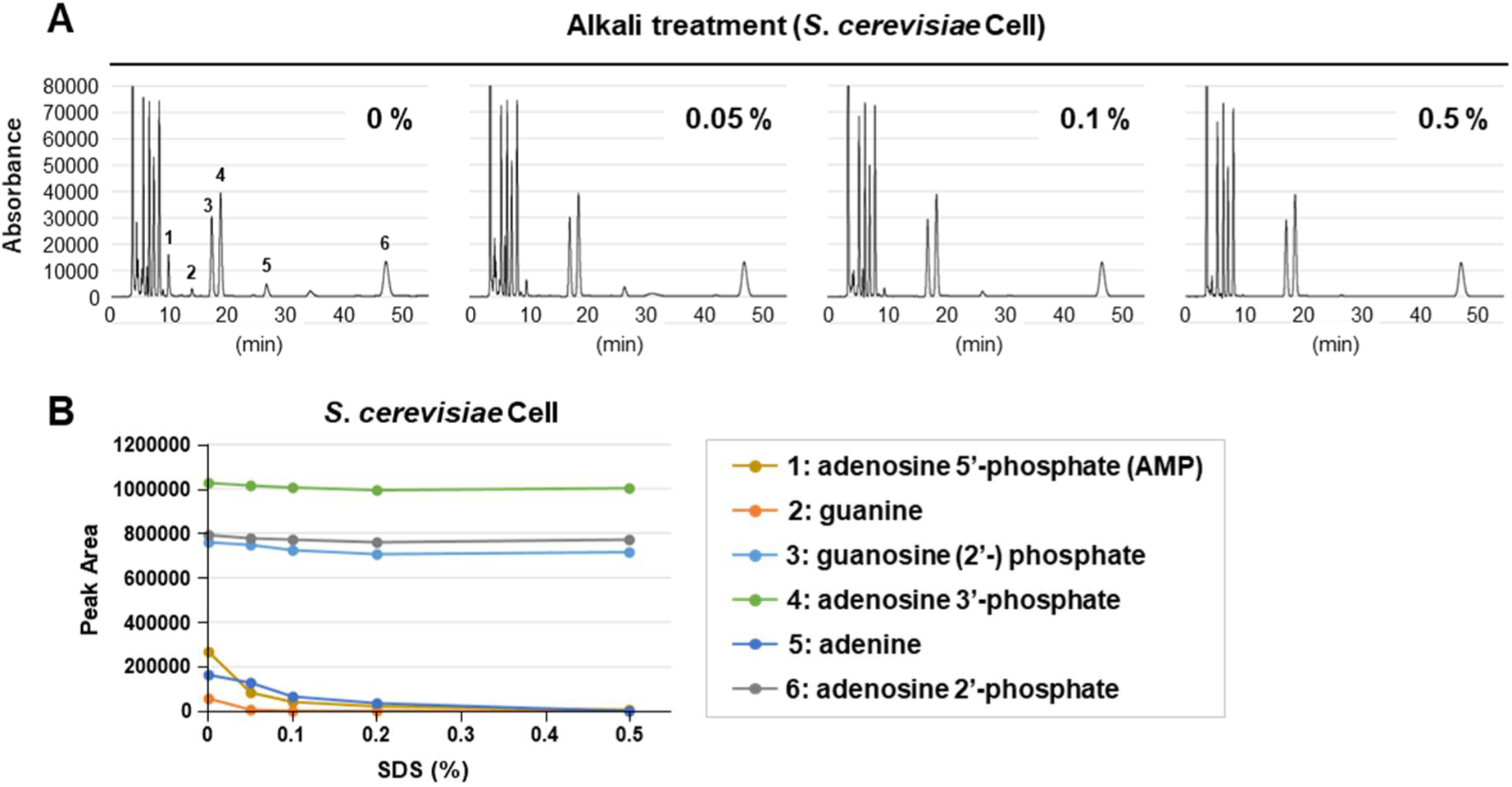
Selective removal of biomolecules by SDS prewashing. (A) Samples prepared from *S*. *cerevisiae* BY4743 cells prewashed with SDS solutions at different concentrations were subjected to alkaline treatment. The samples were analyzed by HPLC, and representative chromatograms are shown. (B) Peak areas of the molecules shown in (A) are plotted across all SDS concentrations (0%, 0.05%, 0.1%, 0.2%, and 0.5%). A representative result is shown.

**Fig. S4.**
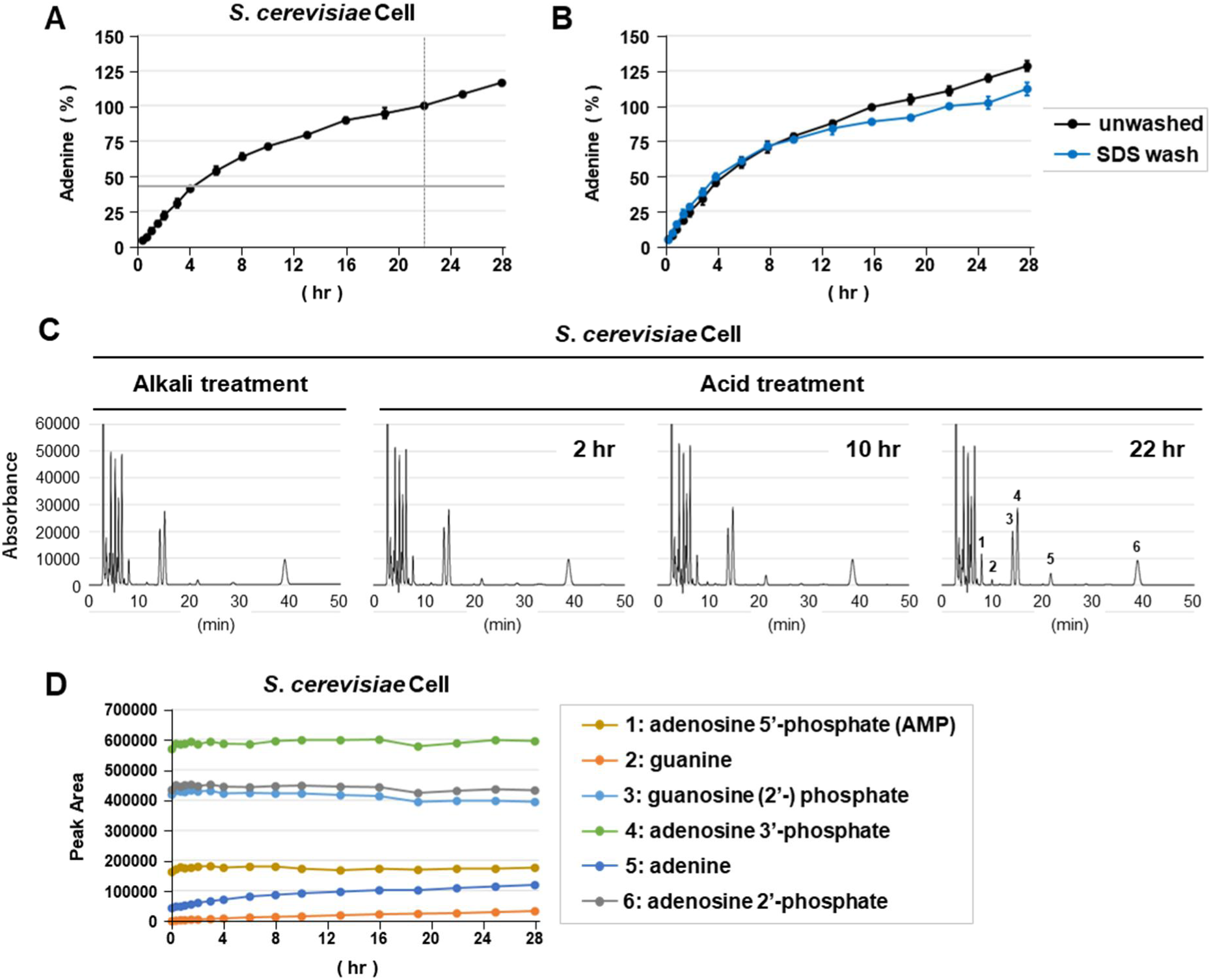
Adenine release kinetics in unwashed *S*. *cerevisiae* cells. (A) Alkali treatment and acid treatment at 35 °C were performed for the indicated time points using unwashed *S. cerevisiae* BY4742 cells (0.3, 0.7, 1, 1.5, 2, 3, 4, 6, 8, 10, 13, 16, 19, 22, 25, and 28 h). The measured adenine amount is shown relative to the 22-h time point (100%). Each measurement was performed three times using independently prepared samples. (B) Except for the absence of SDS prewashing, the treatments were identical to those applied to BY4742 cells shown in Fig. 4A. The adenine amounts in (A) and Fig. 4A are presented relative to the 22-h time point (100%) used in Fig. 4A. (C) Representative chromatograms obtained at selected acid treatment time points in (A) are shown. Each peak is assigned a number corresponding to the molecular species described in Fig. 1. (D) Peak areas of the molecules shown in (C) are plotted across all time points indicated in (A). A representative result is shown (n = 3).

**Fig. S5.**
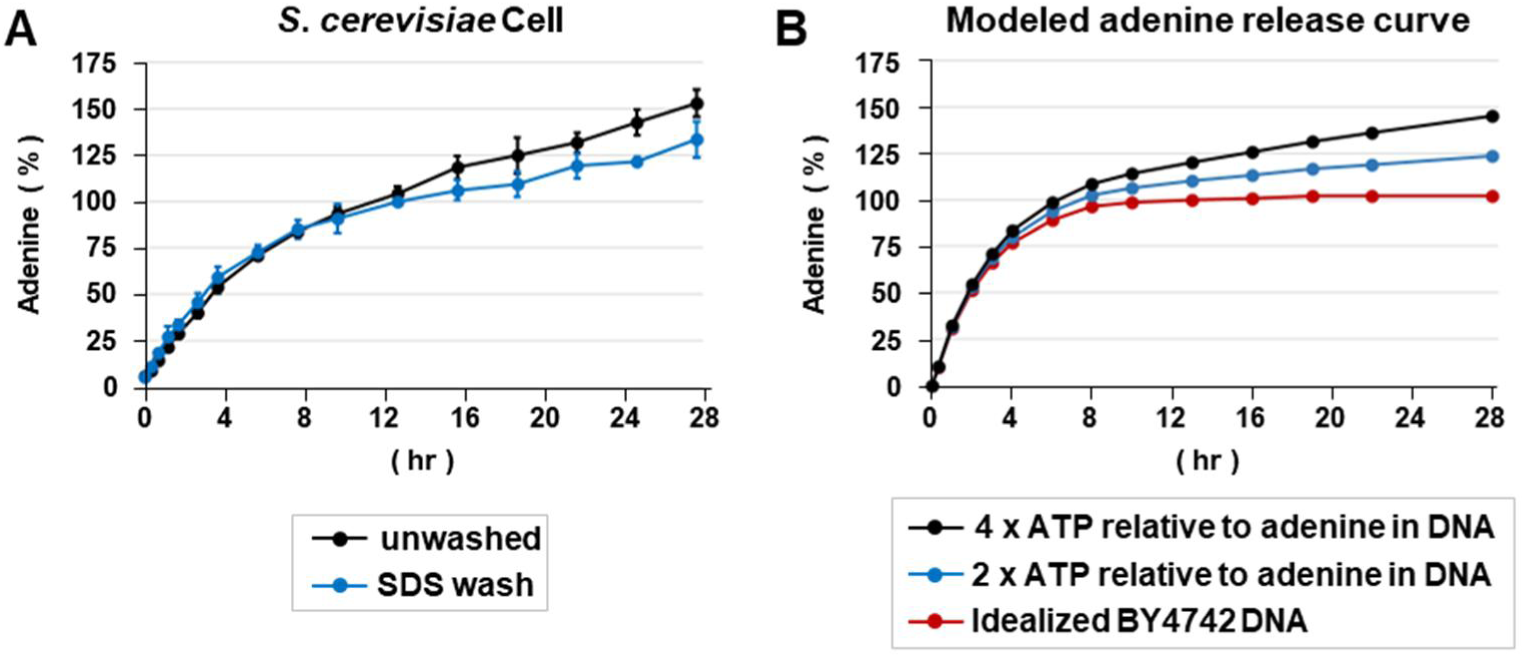
Modeling of adenine release kinetics in SDS-prewashed and unwashed *S*. *cerevisiae* cells. (A) Data from Supplemental Fig. S4 were recalculated and expressed as relative ratios to the value obtained for 13-h SDS-prewashed cells. (B) The adenine release reaction in BY4742 cells was simulated as follows. First, the adenine release curve of purified BY4742 DNA shown in Fig. 4A was idealized and used. Second, the values in Fig. 3B representing the rate of adenine release from ATP at 35 °C were idealized as a simple linear reaction and used. Third, the ATP amount was defined as the peak area ratio in the HPLC chromatogram and set to either twofold or fourfold higher than the amount of adenine derived from DNA. Adenine release was assumed to originate solely from these two components in BY4742 cells for modeling.

**Table S1.** *E*. *coli* and *S*. *cerevisiae* strains used in this study.

| Species | Strain | Genotype | Ploidy | Genome Information |
| --- | --- | --- | --- | --- |
| *Saccharomyces cerevisiae* | BY4741 | *MATa ura3 Δ0 leu2 Δ0 his3 Δ1 met15 Δ0 LYS2* | haploid | JRIS00000000 |
|  | BY4742 | *MAT α ura3 Δ0 leu2 Δ0 his3 Δ1 MET15 lys2 Δ0* | haploid | JRIR00000000 |
|  | BY4743 | *MATa/MAT α ura3 Δ0/ura3 Δ0 leu2 Δ0/leu2 Δ0 his3 Δ1/his3 Δ1 MET15/met15 Δ0 LYS2/lys2 Δ0* | diploid | - |
| *Escherichia coli* | NBRC 3301 | - | - | CP048439, CP048440 |

**Table S2.**
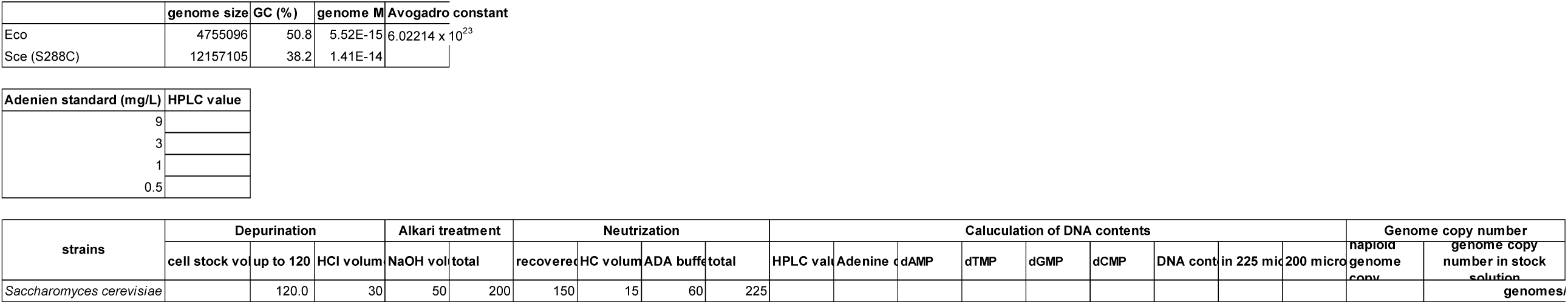
Spreadsheet for calculating genome copy number from adenine amount measured by HPLC.

## Notes

### Competing Interest Statement

The authors have declared no competing interest.

